# Multi-Structure Cortical States Deduced from Intracellular Representations of Fixed Tactile Input Patterns

**DOI:** 10.1101/810770

**Authors:** Johanna Norrlid, Jonas M.D. Enander, Hannes Mogensen, Henrik Jörntell

## Abstract

The brain has a never-ending internal activity, whose spatiotemporal evolution interacts with external inputs to define how we perceive them. We used reproducible touch-related spatiotemporal inputs and recorded intracellularly from rat neocortical neurons to characterise this interaction. The synaptic responses, or the summed input of the networks connected to the neuron, varied greatly to repeated presentations of the same tactile input pattern delivered to the tip of digit 2. Surprisingly, however, these responses sorted into a set of specific response types, unique for each neuron. Further, using a set of eight such tactile input patterns, we found each neuron to exhibit a set of specific response types for each input provided. Response types were not determined by global cortical state, but instead likely depended on the time-varying state of the specific subnetworks connected to each neuron. The fact that some types of responses were recurrent, i.e. more likely than others, indicates that the cortical network had a non-continuous landscape of solutions for these tactile inputs. Therefore, our data suggests that sensory inputs combine with the internal dynamics of the brain networks, thereby causing them to fall into one of multiple possible perceptual attractor states. The neuron-specific instantiations of response types we observed suggest that the subnetworks connected to each neuron represent different components of those attractor states. Our results indicate that the impact of cortical internal states on external inputs is substantially more richly resolvable than previously shown.

**Key points summary:**

- It is known that the internal state of the neocortical network profoundly impacts cortical neuronal responses to sensory input.
- Little is known of how the internal neocortical activity combines with a given sensory input to generate the response.
- We used eight reproducible patterns of skin sensor activation and made intracellular recordings in neocortical neurons to explore the response variations in the specific subnetworks connected to each recorded neuron.
- We found that each neuron exhibited multiple, specific recurring response types to the exact same skin stimulation pattern and that each given stimulation pattern evoked a unique set of response types.
- The findings indicate a multi-structure internal state that combines with peripheral information to define cortical responses; we suggest this mechanism is a prerequisite for the formation of perception (and illusions) and indicates that the cortical networks work according to attractor dynamics.

## Introduction

Behavioural, mental and perceptual functions of the neocortex depend on internal state control. An unresolved issue is to what extent and how internal states affect the responses to a given external input. A state in the brain can be described as the combination of activity in all of its neurons (Spanne & Jorntell, 2015). Since the neurons make synaptic connections with each other, their activities are not independent, which is reflected in reports of constrained ‘realms’ of possible response combinations in populations of neurons (Luczak *et al.*, 2009; Golub *et al.*, 2018). External inputs to the neocortical circuitry, which generate spatiotemporal patterns of activation arising in the multitude of sensors throughout our bodies, further constrain the space of possible neuronal responses (Luczak *et al.*, 2009). An important aspect of perception, and the foundation of illusions, is that a response is determined not only by the quality, or spatiotemporal pattern, of the sensory input but also depends on the current internal state of the cortex (Arieli *et al.*, 1996; Fiser *et al.*, 2004; Curto *et al.*, 2009; Berkes *et al.*, 2011). Although this is by definition a high-dimensional latent state (Stringer *et al.*, 2019b), a more specific embodiment or physiology of the circuitries generating this type of constraint has so far been difficult to identify. This is not surprising given that direct demonstration requires a precise estimate of what the experimental subject is thinking and how that thinking is instantiated in the circuitry, at the time of the stimulus delivery. Otherwise, the internally generated constraints become an uncontrolled variable, which will be interpreted as internal system noise in sensory-evoked responses. Here we aimed to deduce information about the character of the interactions between the time-evolving cortical internal state and external inputs, based on an analysis of the detailed nature of the cortical neuron responses evoked by inputs consisting of several alternative fixed spatiotemporal patterns of tactile sensory activation, each delivered at a high number of repetitions.

It is difficult to achieve exactly reproducible sensor activation patterns in a living organism where the sensors are located in compliant or movable tissue (Hayward *et al.*, 2014) and their exact location or tuning relative to external stimuli are moreover subject to uncontrollable efferent control by the brain. Therefore, we used electrical intracutaneous stimulation to deliver a set of reproducible but also perceptual rich sensory input patterns to the brain (Oddo *et al.*, 2017) (Figure 1A). In order to minimise system noise caused by uncontrollable movements and internal thought processes unrelated to the stimuli (see Methods for further details), and in order to make the rats accept long-term stimulation of the skin electrodes, we used light anaesthesia. The intracellularly recorded signal represents the summed synaptic input from tens of thousands of neurons (out of 25,000,000 neurons in the rat neocortex (Bandeira *et al.*, 2009)). As these neurons, by being connected to the recorded neuron, are by definition part of the same subnetwork, the intracellular signal is a read-out of the instantiation of the current brain state that is specific to the subnetworks connected to that neuron. Here we find that each out of eight pre-defined tactile input patterns generates a wide range of response types that are specific to each neocortical neuron. The findings suggest a multi-component, structured, subnetwork-specific interplay between internal cortical states and sensor input patterns.

**Figure 1.**
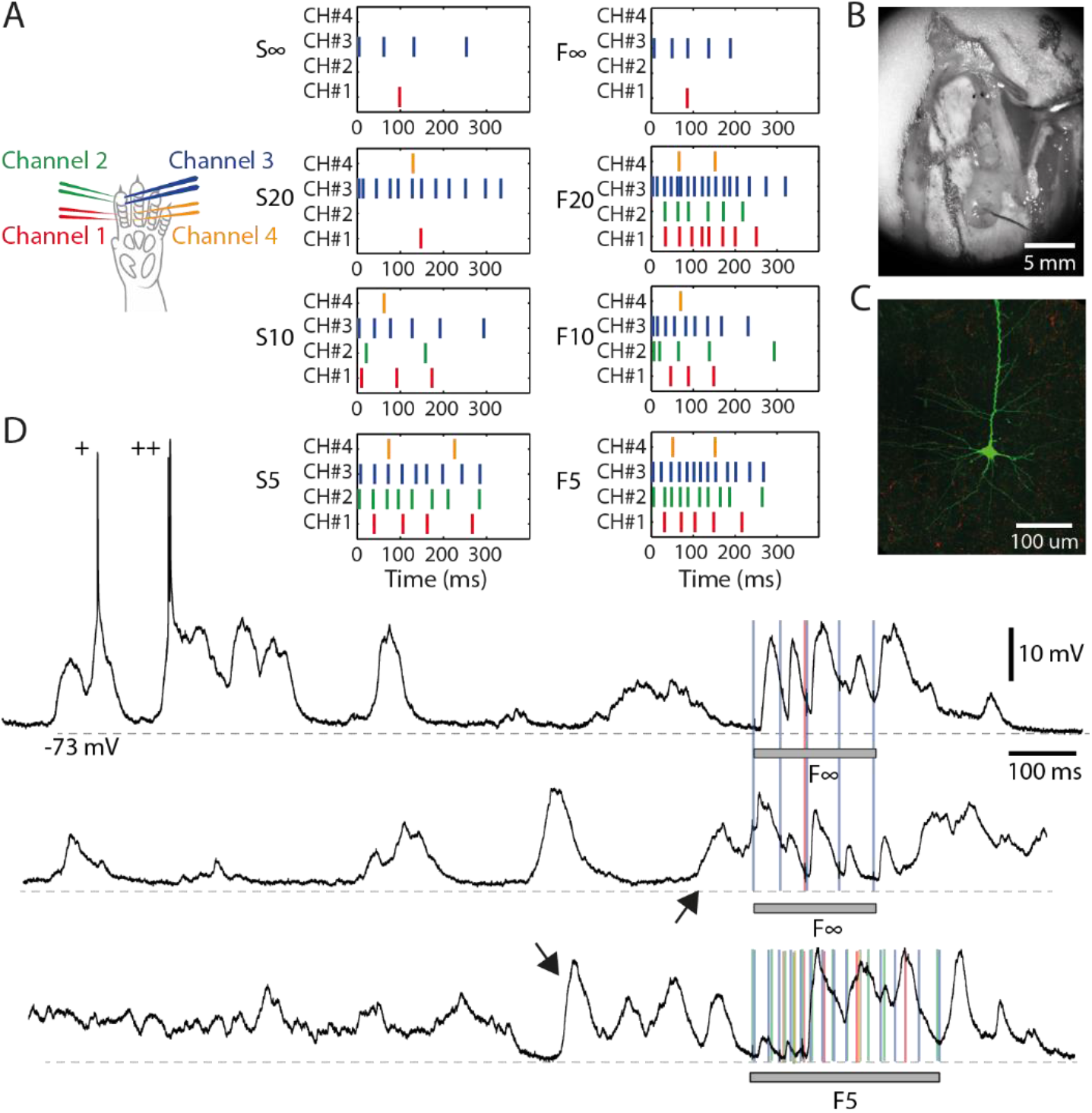
Electrical skin stimulation and general properties of the intracellular responses. (**A**) Locations of the four pairs of intracutaneous needle electrodes (‘channels’) inserted in digit 2 to stimulate the tactile primary afferents. The eight spatiotemporal patterns were re-used from a previous publication (Oddo *et al.*, 2017). (**B**) Neuronal recordings were made in an exposed cortical area of 4 by 2 mm, located in the center of the photo. An electrocorticography (ECoG) surface electrode was placed on a separate, smaller exposed area in the lower half of the photo. (**C**) A stained layer III pyramidal neuron from one of the recordings. (**D**) Intracellular traces illustrating the spontaneous activity and responses evoked by the stimulation patterns (grey horizontal bars). ‘+’ indicates occasional spikes. Arrows indicate specific spontaneous activity patterns. Coloured lines indicate the times of individual stimulation pulses in pattern F(inf) and F5, respectively. Recording from Neuron #7. Stimulus artefacts are truncated for clarity.

## Methods

### Ethical approval

All animal experiment procedures in the present study were in accordance with institutional guidelines and were approved in advance by the Local Ethics Committee of Lund, Sweden (permit ID M118-13 and M13193-2017). All experiments were made using acute preparations under general anaesthesia. All experimental procedures comply with the ‘Principles and standards for reporting animal experiments in The Journal of Physiology’ as defined by the journal.

#### Surgical procedures

Adult male Sprague Dawley rats of male sex (N=10, weight 240–383 g, Taconic) were briefly sedated with isoflurane (3%, 1-2 minutes), anaesthetised with an intraperitoneal injection (40 mg/kg ketamine, 4 mg/kg xylazine) and maintained under anaesthesia with a continuous infusion (ketamine and xylazine in a 20:1 ratio, appx. 5 mg/kg ketamine/hour) administered through an intravenous (IV) catheter inserted into the right femoral vein. A hemicraniectomy (appx 4 by 2 mm) exposed the area of the right somatosensory cortex (Figure 1B). An electrocorticography (ECoG) electrode was positioned on the surface of the brain in order to continuously monitor the depth of anaesthesia by ensuring the presence of sleep spindles mixed with epochs of more desynchronised activity, a characteristic of sleep (Niedermeyer & da Silva, 2005). The level of anaesthesia was additionally characterised by an absence of withdrawal reflexes in response to noxious pinches of the hind paw. The type of anaesthesia used here has little disruptive effect on the physiological network structure at short time spans (in the order of 100s of ms) as judged by the preservation of the order of neuronal recruitment of neocortical neurons in spontaneous brain activity fluctuations (Up states, recordings obtained using multi-electrode arrays in the rat) and stimulus-evoked responses (Luczak & Bartho, 2012). Anaesthesia drags down the overall activity in the neocortical network (Constantinople & Bruno, 2011), though, and in general can be expected to make those networks function with a lower degree of precision. Nevertheless, for the present study, the method of stimulus delivery (see below) would not be accepted by the awake animal and meeting the requirement for long term intracellular recordings was further facilitated by the anaesthesia. To create further mechanical stability, and to protect the brain tissue from dehydration, an agarose gel was applied to cover the exposed part of the cortex. After finishing the neuronal recordings the animal was sacrificed with pentobarbital (140 mg/kg IV).

#### Artificial touch inputs

In order to achieve as realistic spatiotemporal patterns of tactile sensor activation as possible, while preserving the aim of high reproducibility of the patterns, we used an artificial fingertip equipped with four neuromorphic sensors to generate a set of spatiotemporal patterns of skin activation to be used in the electrical interface with the rat skin. The procedures have previously been described in greater detail and the patterns used here were the same as before (Oddo *et al.*, 2017; Genna *et al.*, 2018; Enander *et al.*, 2019). As discussed in this previous work, the artificial fingertip allowed synthesising spatiotemporal patterns of skin sensor activation at quasi-natural firing rates that follow a natural overall temporal modulation, or ‘envelope’, that the biological skin sensors are known to display under dynamic indentation (Jenmalm *et al.*, 2003). This aspect is important because the circuitry of the cortex can be expected to have experienced many events with similar envelopes of tactile afferent activity and is therefore likely to have adapted its circuitry structure to effectively process variations of that overall pattern of activity modulation (Berkes *et al.*, 2011; Oddo *et al.*, 2017). By delivering this input electrically to the primary afferents in local skin sites we by-passed the step of potentially variable skin sensor activation that occur even with highly controlled mechanical skin stimulation, so that the decoding capacity of cortical neurons could be studied in relative isolation. Just like natural tactile inputs, the input we provided can be expected to be distributed and processed through the neuronal networks in the cuneate nucleus, thalamus and neocortical circuitry before it reaches the neurons we recorded from. Hence, the measured responses are bound to reflect at least in part the natural processing mechanisms of the brain. Accordingly, in humans, electrical nerve stimulation with a much lower resolution than in the present set of experiments are known to generate sensory impressions that are in part perceived as unnatural but also to generate diversified and meaningful tactile percepts (Tan *et al.*, 2014; Oddo *et al.*, 2016).

The spatiotemporal tactile input patterns were generated as follows: the bionic fingertip was moved against different types of objects and the number label of each pattern (Figure 1A) indicates the radius of the curvature of different objects used to generate each specific pattern, whereas the letter of the label indicates the adaptive tuning of the biomorphic sensors (S, F; for slow and fast, respectively). These spatiotemporal stimulation patterns were delivered via four pairs (or channels) of intracutaneous needle electrodes (Figure 1A) in a pre-defined random order, where each pattern lasted for less than 340 ms and the consecutive deliveries of the patterns was separated by 1.8 s in order to allow a relaxation of the cortical activity between consecutive deliveries of stimulation patterns (Oddo *et al.*, 2017). Each of the eight patterns was delivered 100 times, except for four out of the total thirteen neurons for which the recording was lost after 36, 47, 50 and 80 repetitions, respectively. In addition, for each of the four stimulation channels used, we delivered the same number (up to 100) of repetitions of isolated single-pulse stimulations. These isolated single-pulse stimulations were delivered in chunks of five stimulations from the same channel separated by 300 ms from each other. Thus, for each channel there was 20 such chunks intermixed with the stimulation patterns in a random order. Each channel was stimulated at 0.5 mA with a pulse width of 0.14 ms, which is higher than the threshold of about 0.2 mA reported for tactile afferents using this type of electrocutaneous stimulation (Rasmusson & Northgrave, 1997; Bengtsson *et al.*, 2013), but lower than the threshold for activating nociceptive afferents (Ekerot *et al.*, 1987).

#### Neural recordings

We made recordings in the region of the primary somatosensory cortex (S1) of the forepaw, as estimated by the focus of the local field potentials (measured between layers III and V, corresponding to the depths of maximum field potential negativity recorded in each track). The coordinates of this region were −1.0–1.0 mm relative to bregma and 3.0-5.0 mm lateral to the midline (Figure 1B). Individual neurons were recorded with patch clamp pipettes in the intracellular, whole cell current clamp mode. Patch clamp pipettes were pulled from borosilicate glass capillaries to 6–15 MOhm using a Sutter Instruments (Novato, CA) P-97 horizontal puller. The composition of the electrolyte solution in the patch pipettes was (in mM) potassium-gluconate (135), HEPES (10), KCl (6.0), Mg-ATP (2), EGTA (10). The solution was titrated to 7.35–7.40 using 1 M KOH. During slow advancement of the recording electrode (approximately μm per second) made with positive pressure applied, electrode tip resistance and responses evoked by electrical skin stimulation were continuously monitored to identify encounters with neurons. Once encountered, the positive pressure was changed to a negative pressure, and a weak hyperpolarizing current was applied with the aim of obtaining a GigaOhm seal on the neuron. Successful access to the intracellular signal of the neuron, following additional negative pressure once the seal was established, was followed by a release of pressure and the start of the data collection. Using a weak hyperpolarizing bias current, neurons were prevented from spiking. All intracellular data was digitised at 100 kHz using CED 1401 mk2 hardware and Spike2 software (Cambridge Electronics Devices, CED, Cambridge, UK). The criteria used for inclusion of an intracellular recording, or the time period of such a recording to be included in the analysis, were a stable membrane potential of <−55 mV in Down states, a spike amplitude of >25 mV before and after the termination of the protocol and a peak-to-peak difference between the Up and Down states of >10 mV. All neurons recorded were putatively located within layer III-V based on the recording depth measured from the cortical surface (Narayanan *et al.*, 2017). For identification of neuron identity, in addition to depth, we used the nature of the firing during spontaneous activity (i.e. if the neuron was fast-spiking, bursting and what duration and intensity of bursts the neuron displayed). All neurons recorded here exhibited infrequent bursts of two or three spikes but had an absence of longer bursts or sustained periods of high firing. Based on this criterion, they were considered to be pyramidal cells rather than interneurons (Luczak *et al.*, 2009). Three out of the thirteen successfully recorded neurons were also stained with neurobiotin and histologically recovered. They were thereby confirmed to be pyramidal neurons located in layer III (Figure 1C).

### Quantification and statistical analysis

#### Post processing - general

The neuronal recording signal was imported from Spike2 to Matlab (2016a, Mathworks), where it was low-pass filtered using a moving average over 50 μs, i.e., 5 samples width given a 100 kHz sampling rate. Stimulation artefacts were removed using a combination of adaptive filtering and blanking of artefacts. Occasional remaining intracellular spikes were removed using adaptive filtering, with a recursive fitting algorithm that created a generic spike shape for the neuron (Mogensen *et al.*, 2019), which could be subtracted from the signal at all occurrences of the spike. This allowed for excitatory postsynaptic potential (EPSP)-like events to be detected also when the membrane potential was influenced by spiking activity. Since all evoked responses were analysed by visual inspection, there was a quality check of the intracellular signal throughout the recording period.

#### Post processing – clustering algorithm for separation of response types

On visual inspection, the time-voltage curves of the intracellular responses to repetitions of a given stimulation pattern appeared to fall into certain classes, or types, where the differences between responses of different types appeared to be larger than the variability of different responses within any given type (Figure 2A).

**Figure 2.**
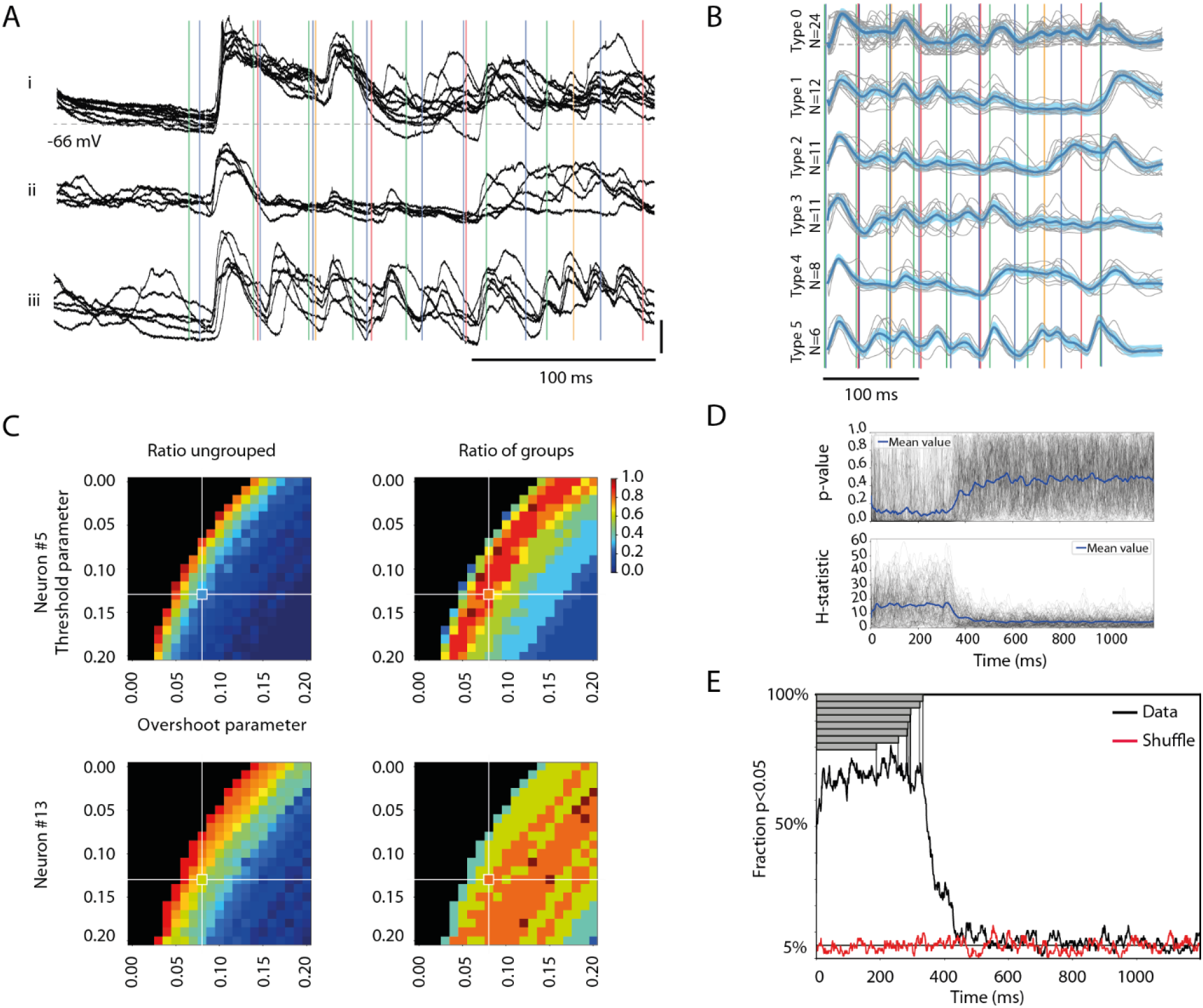
Different response types evoked by the same stimulation pattern. (**A**) Each panel i-iii shows, for a sample cell (Neuron #5), examples of qualitatively different superimposed raw data traces, evoked by the same stimulation pattern (S5). Note that the stimulation pattern (vertical coloured lines) outlasted the traces. Stimulus artefacts are reduced for clarity. (**B**) The six identified response types for all responses evoked by the 100 repetitions of this stimulation pattern in this cell are shown separately. The mean response of each type is indicated by the thick blue curve, the threshold difference by the light blue span and the raw traces are indicated in grey, alongside the stimulation pattern. (**C**) Sensitivity analysis of the parameter settings (for ‘Threshold’ and ‘Overshoot’, respectively, see Methods) of our response type classification/clustering method illustrated for two different neurons. The impacts of the parameter settings on the fraction of responses ending up as ungrouped (ratio ungrouped) and on the number of response types identified versus the maximal number of response types identifiable within the parameter space (ratio grouped) are illustrated for both sample neurons (Neuron #5 is also illustrated in panels **A-B**). Note that the parameter settings used for the main analysis are indicated by the cross-hairs. (**D**) The Kruskal Wallis test result of the separation between the members of each response type from the members of other response types evoked by the same stimulation pattern in the same neuron, plotted as a separate time curve for each stimulation pattern (N=104) (grey traces). The blue trace represents the average across all stimulation patterns. The corresponding plot for the H statistic is shown in the diagram below. (**E**) Time evolution of the specificity of the responses of each response type. The black curve is the fraction of all response types where specificity could be detected (at p<0.05, Kruskal-Wallis test). The red curve shows the corresponding fraction for responses with shuffled response type labels (within each stimulation pattern, for each cell). Grey bars show the durations of the eight stimulation patterns.

To quantitatively evaluate this potential clustering, we used an unsupervised clustering method. This clustering method was merely based on the amplitudes of the intracellular membrane potential response across the whole series of time windows that constituted the response. Thereby, it is based on clear neurophysiological metrics and in this respect differs from traditional machine learning algorithms, in which it is typically not clear which metrics underlie the clustering.

The clustering method defined both the number of response types and the number of members/individual responses that belonged to each response type. The clustering method also had to be capable of reporting zero response types, if no response types were possible to identify according to the criteria of the clustering method. Furthermore, the uniqueness of the time-voltage curve for each response type had to be evaluated against all other responses as a validation of that the grouping into response types was sound. The following procedure was used to sort the intracellular response curves into types:

1. For each cell, the 350 ms time-voltage curves from the onset of stimulation for each of the (up to) 100 repetitions of a given stimulation pattern were compared.
2. To remove high-frequency fluctuations the responses were low-pass filtered (with a 1 ms wide moving average) and re-sampled to 1000 Hz. In order to focus on the temporal shape of the responses, a moving average of 100 ms was subtracted from the signal to remove its offset and the response voltage was then normalised to a 1.0 - 0.0 range based on the highest peaks and deepest troughs for each 350 ms time-voltage curve. This was made to ensure that the method captured the shape of the response curve over time.
3. For each neuron/stimulation pattern, each time-voltage curve was subjected to pairwise comparisons against all other time-voltage curves evoked by the same stimulation pattern in the same neuron. The evaluation of similarity between each pair of time-voltage curves was done on a per sample time point basis (i.e. 350 sample points per time-voltage curve), where the difference for each sample point was calculated. If the difference for a sample point was below a threshold value (this ‘Threshold difference’ was set to 0.13 normalised units, see Figure 2C for the sensitivity analysis of how this threshold was chosen), the difference was set to zero for that sample point. If the difference was above the threshold value, the overshoot was calculated as the absolute value of the actual difference above the threshold value for that sample point. If the mean overshoot across the 350 samples fell below a threshold (this ‘Overshoot threshold’ was set to 0.08 normalised units, see Figure 2C) the two time-voltage curves were classified as the same response type. This procedure helped reducing sensitivity to high-amplitude transient membrane potential shifts, while emphasising persistent low amplitude differences between time-voltage curves in the classification/clustering.
4. This procedure was repeated so that all time-voltage curves were compared with all others of the neuron/stimulation pattern (i.e. typically 100 responses). This resulted in the identification of several types of responses. The members of the response type with the largest number of members were removed from the set of 100 responses, and the sorting (steps 1-4) was repeated until there were no remaining responses left to sort.
5. Last, for the set of detected response types, for each response type with less than 5 members, the members were categorised as belonging to the ‘ungrouped’ response type and that response type was no longer a valid response type.

#### Sensitivity analysis for the selection of parameter values for the clustering method

The clustering method described above is unsupervised, but depend on the choice of parameter values for the Threshold difference and the Overshoot threshold (step 3 above). The Threshold difference parameter is an absolute voltage distance to the reference response on a per sample point basis (Figure 2B, light blue area). The Overshoot threshold is the mean of that distance across all sample points.

To test the sensitivity of the method to the choice of parameter values, we explored the resulting outcome (number of response types identified, fraction of the responses that fell into the ungrouped category and the accuracy by which the identified response types could be identified) across parts of the parameter space for two neurons’ sets of responses to different stimulation patterns. This exploration is visualised in Figure 2C and 3B, which suggested that our choices of parameter values were located in the middle of a smooth landscape of outcomes.

**Figure 3.**
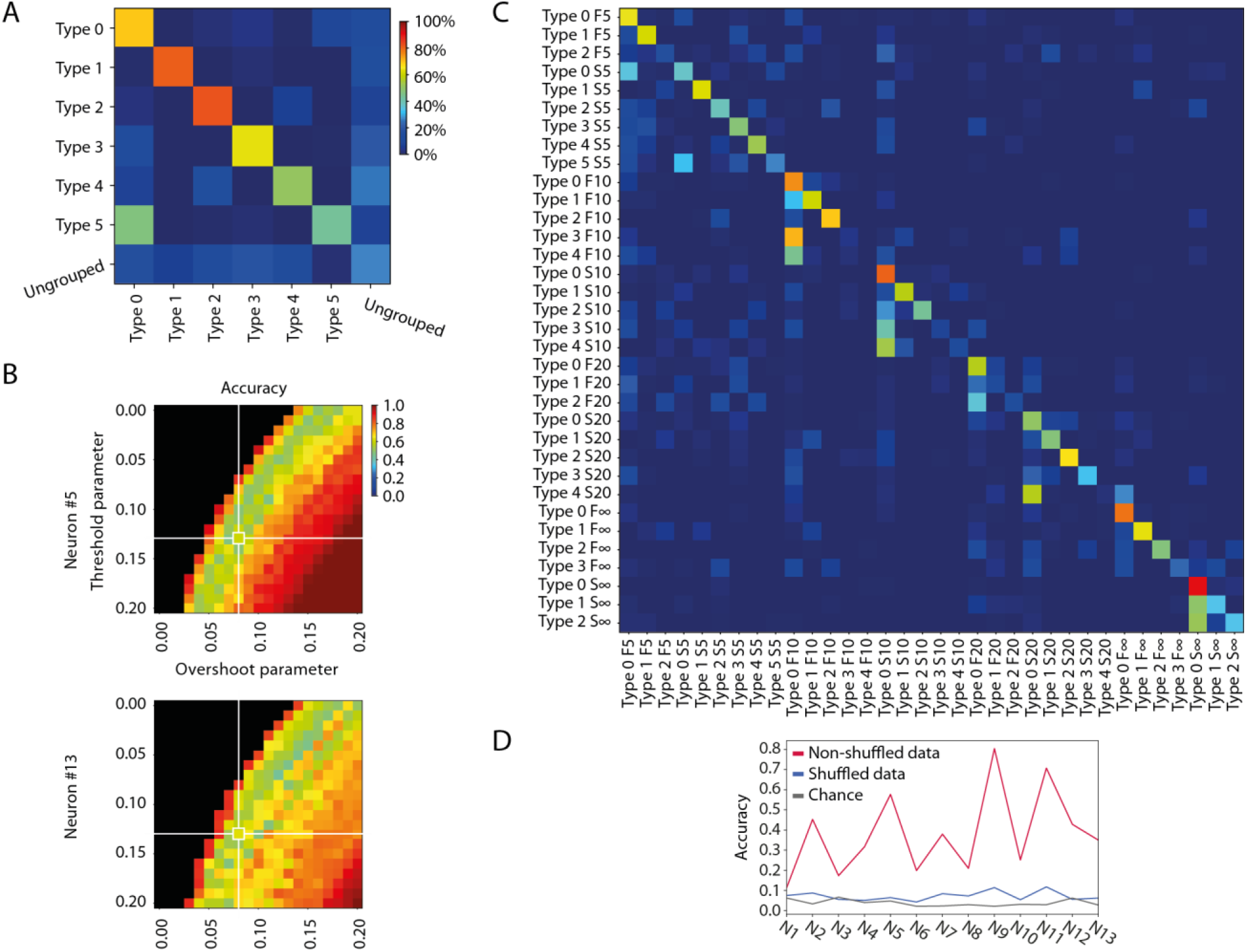
Specificity of the different response types evoked by the same stimulation patterns in the same neuron. (**A**) Confusion matrix for the members of each identified response type for the sample stimulation pattern and neuron illustrated in Figure 2A-B. Decoding accuracy for each response type is indicated by the colour code indicated in the calibration bar. (**B**) Sensitivity analysis of the parameter settings of our response type clustering method illustrated for two different neurons, here with respect to the average response type decoding accuracy (see A) but across all stimulation patterns. (**C**) Confusion matrix for all response types identified for all stimulation patterns for this sample neuron. Same colour code as in A. (**D)** To illustrate the specificity of the response types recorded across all eight stimulation patterns, this plot indicates the decoding accuracy across all stimulation patterns for each neuron. The figure also displays the chance level for each neuron, and the decoding accuracy when the responses recorded for each neuron were shuffled.

#### Statistical evaluation of the identified response types

We first tested whether the response types of each simulation pattern, as identified by the clustering method above, were different from each other. For each sample point (350 sample points per response, see step 2 above), we tested if the distributions of the amplitudes in each cluster/response type were different from each other using Kruskal-Wallis test. For each set of response types for each stimulation pattern, this procedure was repeated for each sample point up to 1200 ms after the onset of the stimulation pattern, i.e. for a much longer time than the duration of the stimulation pattern, which yielded a time curve of the p-value that could be visualised individually for each of the 104 stimulation patterns (8 patterns for 13 neurons) in the whole data set (Figure 2D). By defining a threshold p-value of <0.05, we could also display the results of this analysis as a time curve of the fraction of the response types fulfilling this criterion (Figure 2E). In the latter analysis, we also tested whether a shuffling of the response type labels between all the responses evoked by a specific stimulation pattern in a specific neuron resulted in a collapse of this fraction down to the 5% level, which is the expected value for randomly distributed responses (Figure 2E). This shuffled analysis also functioned as a control for the multiple comparisons problem.

#### Evaluation of the specificity of the identified response types for each stimulation pattern in the same neuron

We also evaluated the separability of the whole membrane voltage time curve vectors for each of the identified response types, for each stimulation pattern in each neuron separately, as well as their separability versus the ‘ungrouped’ responses, using a combination of PCA and kNN-classification (Figure 3A).

We started by using the mean membrane voltage time curve vector of each response type (see step 2 above) to calculate the Principal Components (PCs). The number of PCs used was the number required to explain at least 95% of the total variance of the mean signals. Finally, we used the principal component coefficients to transform each individual recorded membrane voltage time curve vector from the time domain to the principal component domain, reducing the dimensionality of each response from M = 350 (sample time points) to N = [1-6] (PCs).

The position of each response in PC space was then used for verification of the accuracy of the classification/type identity of each response. In order to determine the separability of the detected response types, and their possible confusion with the ‘ungrouped’ responses, we used a kNN classification procedure. Half of the responses were randomly selected as the training set. For each response belonging to the test set we identified the 4 closest responses in the training set by calculating the Euclidian distance in PC space. The response was classified as belonging to the same response type as the relative majority of the 4 neighbours, where the classification either matched the response type assigned to it by the clustering method, or not. We performed 40 iterations of the classification, each with a different training set, and averaged the fraction of correctly classified responses in each iteration to get the mean correct classification value for each response type. The results of the ensuing kNN decoding are visualised in confusion matrices, which were also used to extract the mean decoding accuracy and the F1 score.

#### Evaluation of the specificity of the response types across all stimulation patterns in the same neuron

To evaluate if the identified response types were specific to the stimulation pattern, we again used PCA and kNN-classification. Here, the PCs were calculated based on the mean membrane voltage time curve vector of each response type across all eight stimulation patterns. Because the total number of responses considered in this case was in the order of 800 rather than 100 (as above), the dimensionality of the responses was reduced to N = [8-40] PCs, rather than [1-6] PCs. Also, the number of neighbours evaluated for the kNN classification were in this case nine (N=9).

#### Evaluation of the specificity of the response types across the same stimulation pattern in different neurons

We also used PCA and kNN decoding to evaluate if the response types were specific to the neuron, for each specific stimulation pattern (see for example Figure 4B), using the same type of approach as above. Here, the PCs were calculated based on the mean membrane voltage time curve vector of each response type, for a given stimulation pattern, across all thirteen neurons.

**Figure 4.**
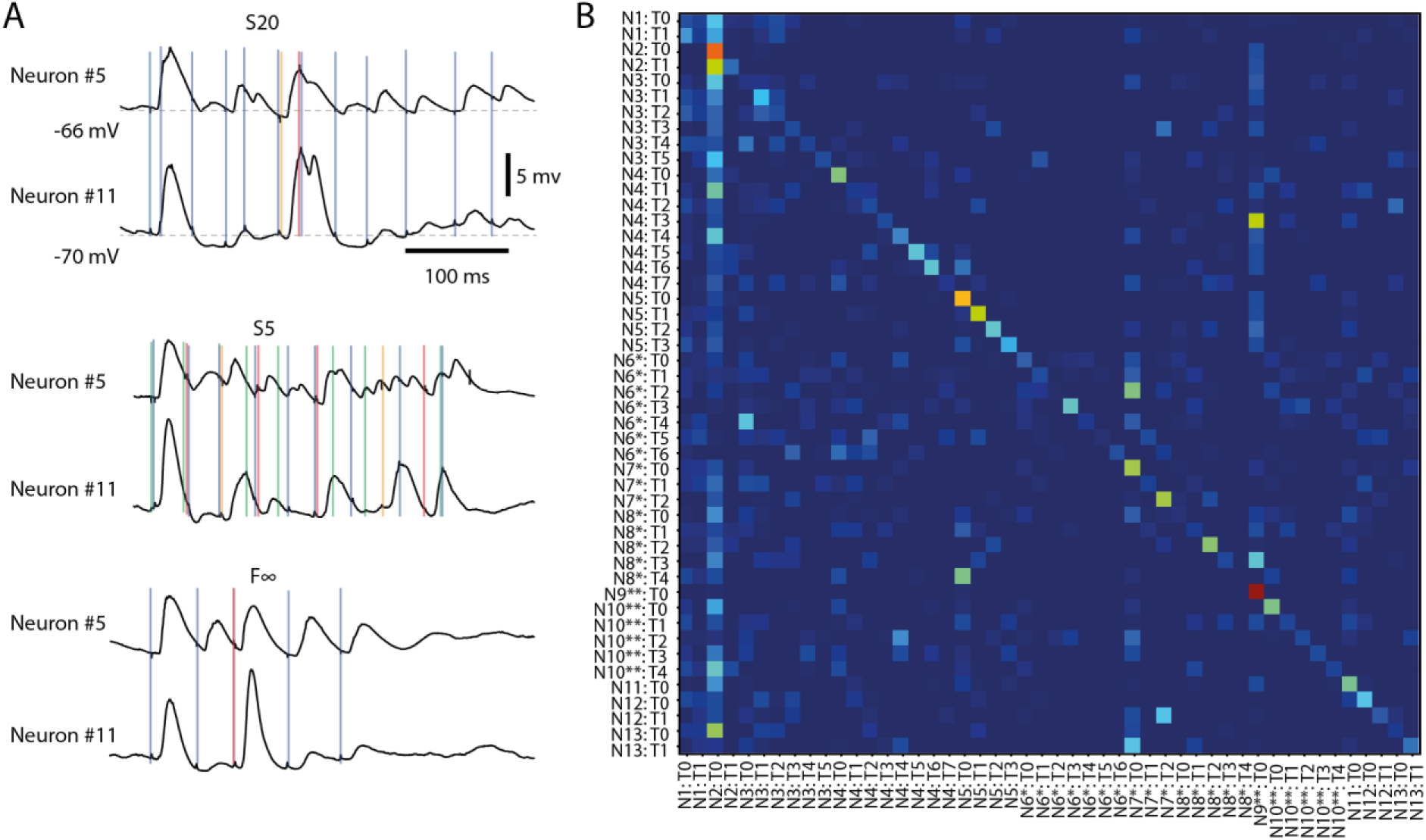
Different response types evoked by the same stimulation pattern in different neurons. (**A**) Averages of intracellular responses of neuron #5 (N=100 responses) and neuron #11 (N=80 responses) to stimulation pattern S20 (top). Vertical coloured lines indicate the individual stimulation pulses of that stimulation pattern. Similar display for the average responses to two other input patterns (middle, bottom). The responses of Neuron #5 to pattern S5 are also illustrated in Figures 2A-B and 3A. (**B**) Confusion matrix for all response types identified for all neurons for stimulation pattern F∞. Neurons (N#) recorded in the same experiments are indicated by * and **, respectively. T# indicates the response type.

#### Brain state segmentation

During each neuronal recording, a parallel ECoG signal was recorded at a sample rate of 1 kHz from the surface electrode placed on the surface of the cortex nearby the recording site (Figure 1B). To segment the recording into epochs occurring during synchronised versus desynchronised ECoG states, as previously shown (Enander *et al.*, 2019), the spectral density of the ECoG was calculated with a segment length of 1,000 ms, an overlap of 125 ms and a constant (mean) detrending. The spectral density of Delta, Theta and Alpha bands (0–12 Hz) was summed for each segment and the compound value was used for the remainder of the analysis. A desynchronised segment of ECoG was assumed to occur when the compound spectral density dropped below the compound spectral density median for at least two segments in sequence. Therefore, at the onset of every stimulus presentation, and thereby for each response type, there was an identified ECoG state. The fraction of stimulus presentations that occurred within the desynchronised state relative to the total number of presentations was used for statistical comparisons (Figure 5). As the probability of a brain state is expected to influence the observed ratios of occurrences of events within each brain state the statistical method used to make the comparisons was the paired t-test.

**Figure 5.**
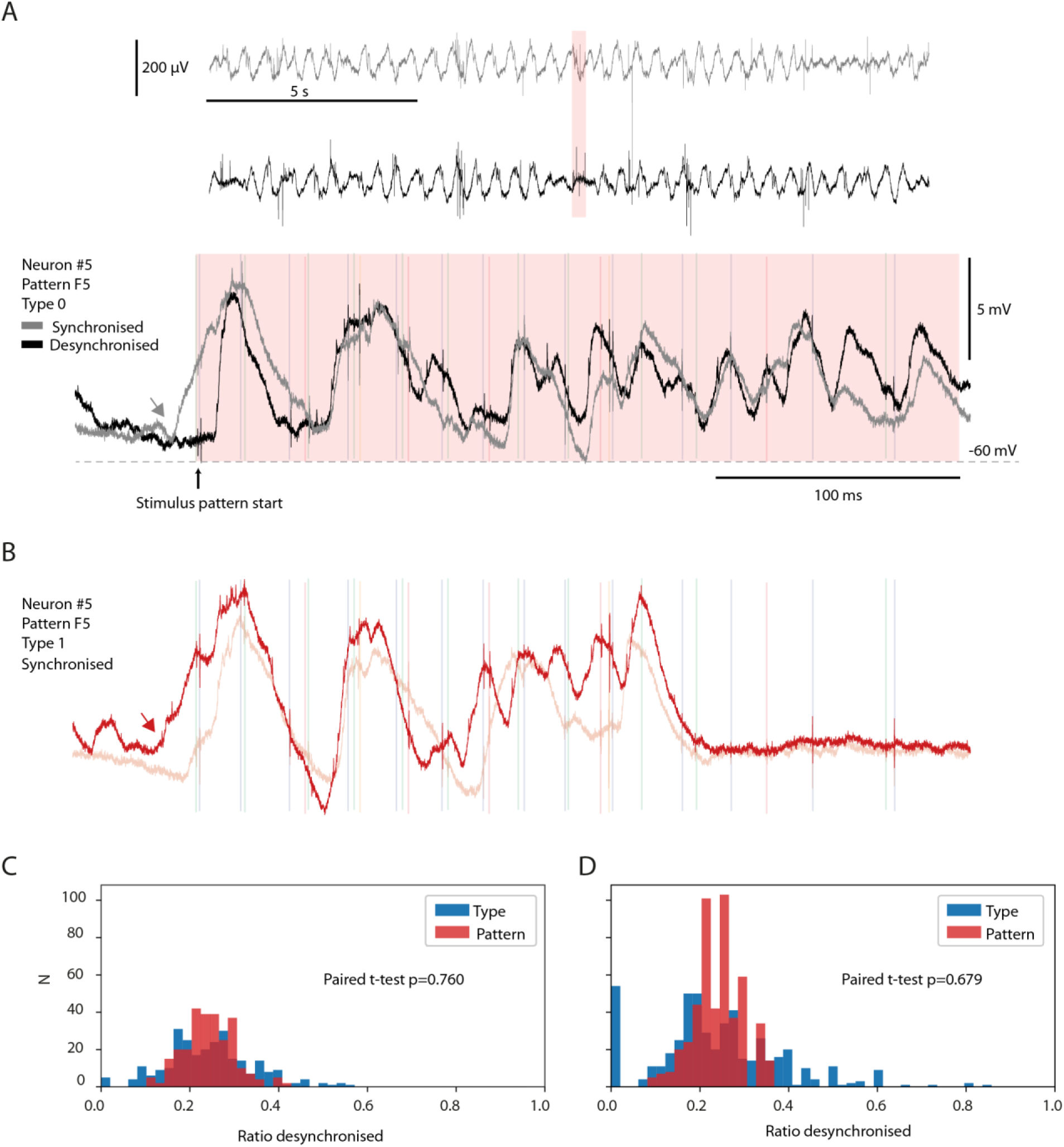
No preferred ECoG state for specific response types. (**A**) Two sample responses belonging to the same response type evoked during synchronised and desynchronised ECoG states. Top two traces are time-compressed ECoG recordings. The pink time zone is expanded below to illustrate the corresponding intracellular responses of a recorded neuron, superimposed. Note the rising Up state starting before the onset of the stimulation (grey arrow) for the grey trace (synchronised). (**B**) Two sample responses from the same neuron and stimulation pattern as in (A), but from response Type 1. Both traces occurred during synchronised ECoG states. The red trace had a rising Up state that started (red arrow) at almost the exact same time as in the grey trace in (A). (**C, D**) The fraction of responses occurring in the desynchronised ECoG state for the responses evoked by each stimulation pattern is shown in red, for each neuron and for each member of a response type (blue). The fraction for each response type is hence paired with the corresponding fraction for the stimulation pattern. The bar chart in B illustrates the results of this investigation only for response types with more than nine members (nine responses). In D, the same display but for all response types identified (including those with less than 10 members, N=494).

#### Post-processing – responses evoked by isolated single-pulse stimulation pulses

As described above, isolated single-pulse stimulations were delivered for each stimulation channel separately in sequences of five repetitions separated from each other by 0.3 s, whereas different such chunks of five pulse stimulations (N=20 per channel) were randomly intermixed with the full stimulation patterns. We analysed the responses to such isolated single-pulse stimulations with respect to the onset latency time, the time-to-peak and the response amplitude, using a tailor-made point-and-click user interface (Figure 6, Table 2).

**Figure 6.**
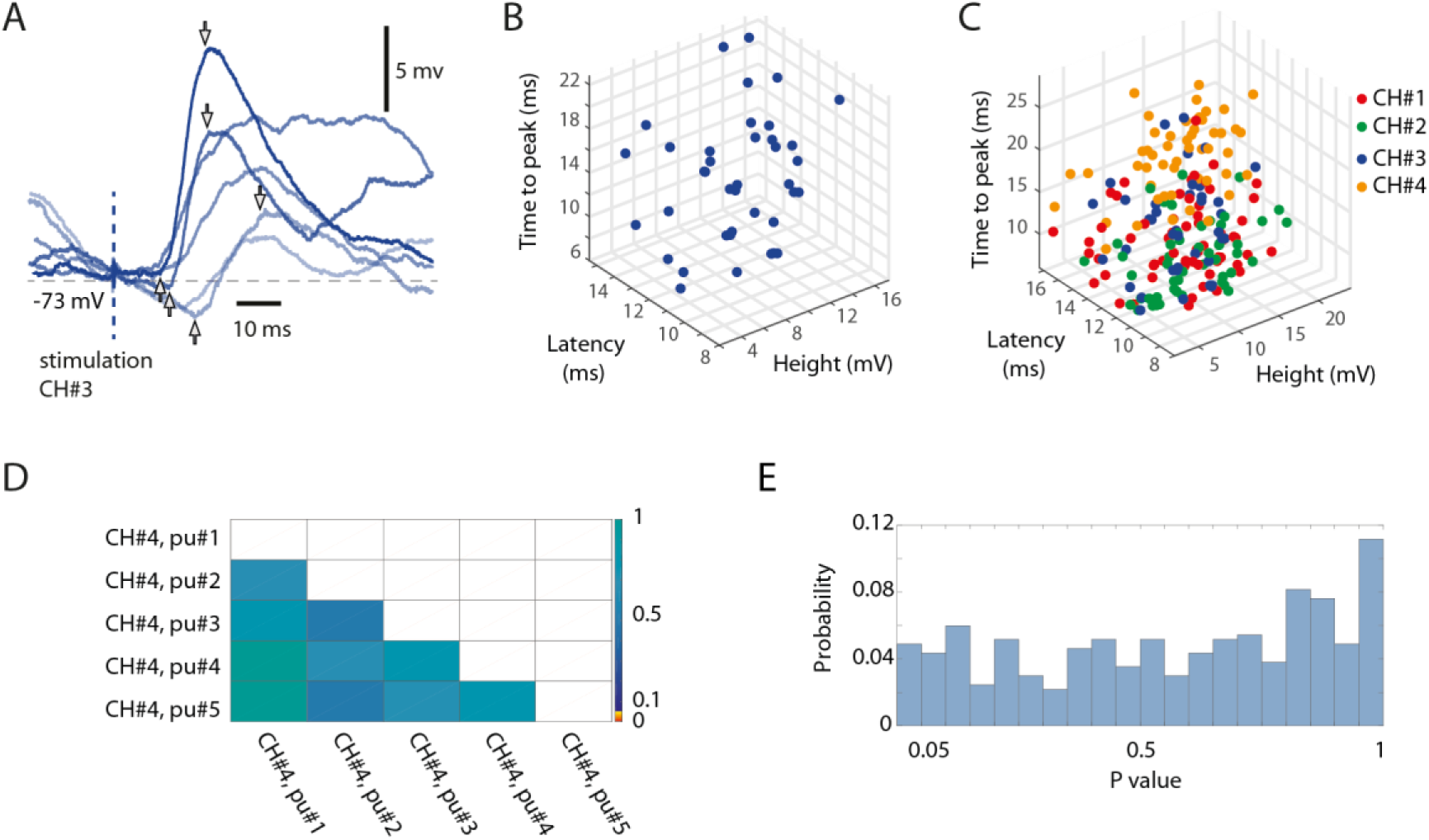
Random variability of the isolated single-pulse responses. (**A**) Six superimposed raw intracellular traces evoked by isolated single-pulse stimulations to channel #3, recording from Neuron #7. Pairs of arrows indicate the response onset latency time, the time-to-peak and the peak amplitude for three sample responses. (**B**) 3D plot of the measured parameters for all responses evoked by isolated single-pulse stimulation of channel #3 in the same neuron. (**C**) Similar display for all four channels used. (**D**) The matrix presents the outcome of a statistical investigation of whether the rank order of the single-pulse stimulations within the 5 pulse chunks (each pulse separated by an interval of 0.3 s) had a statistically significant impact on the response amplitude, by comparing the 20 responses of any one group (the first pulse, for example) to those of each other group (the third pulse, for example) in turn (Wilcoxon rank sum test for pairwise comparisons, p-values are indicated according to the colour scale). (**E**) For the investigation of which an example is shown in D, this histogram summarises the result (the distribution of p values) across all stimulation channels and all neurons (binned).

#### Post-processing – responses evoked by individual stimulation pulses within patterns

We also performed an analysis of the responses to the individual pulses within the different stimulation patterns (Figure 7). We constrained the analysis to the following set of individual stimulation pulses: Each of the eight spatiotemporal stimulation patterns consisted of 5-33 individual stimulation pulses and a total sum of 152 pulses altogether in the eight patterns used (Figure 1A). The time between subsequent stimulation pulses within the stimulation patterns varied in the span 1-123 ms. This means that in some cases, the intervals between subsequent stimulation pulses were too short to identify which of the pulses generated the recorded response. Since the scope of this part of the analysis was to investigate the response to specific in-pattern stimulation pulses, only stimulation pulses that were temporally segregated from previous and subsequent pulses by at least 10 ms were included (as the average response latency time was 11 ms, Table 2). Based on this selection criterion, 52 out of the total 152 pulses were included in the analysis of the responses evoked by the individual stimulation pulses within stimulation patterns.

**Figure 7.**
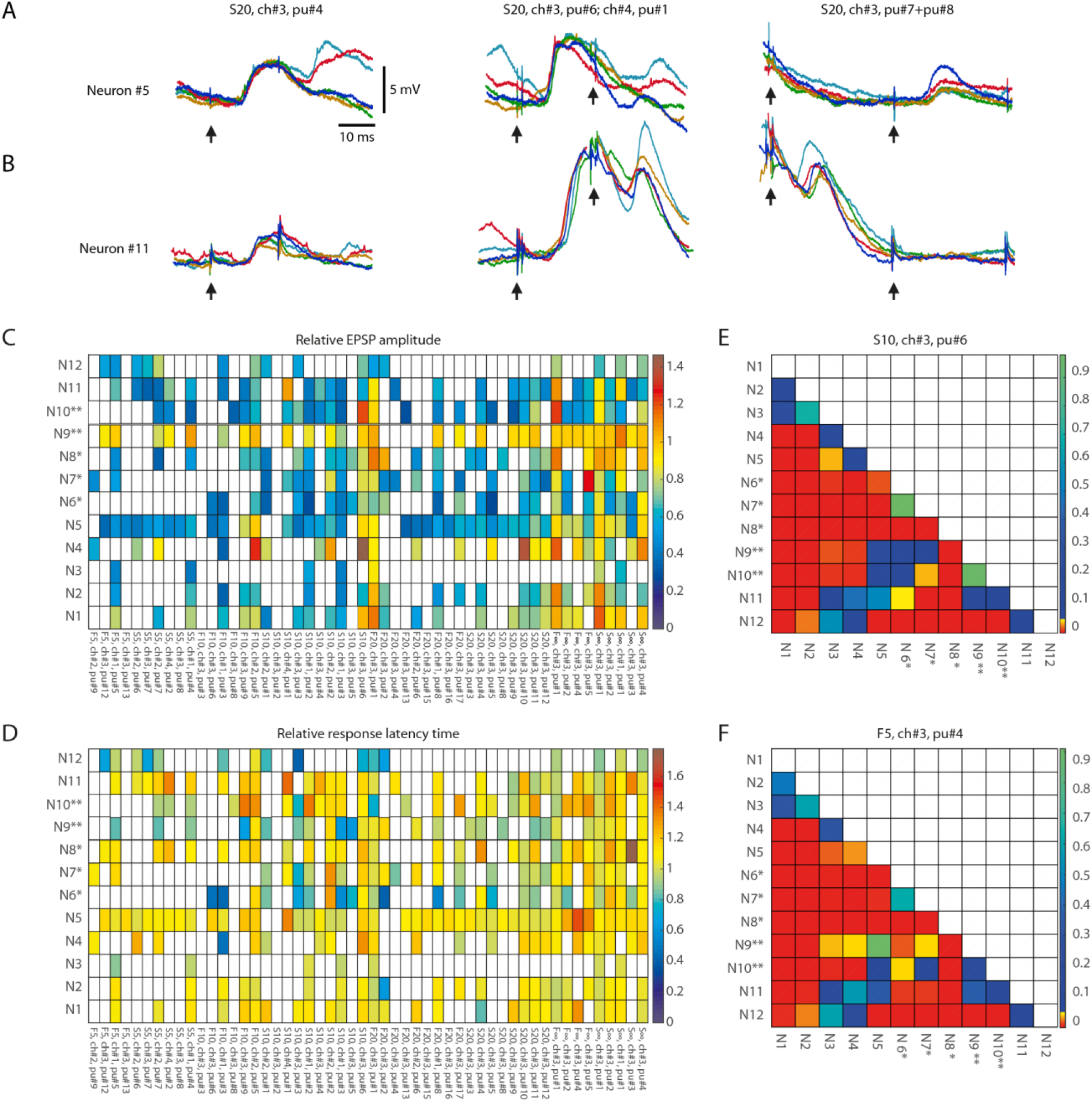
Response metrics differences between neurons for isolated within-pattern stimulation pulses. (**A, B**) Examples of the variation of the responses to single-pulse stimulation of the same channel depending on the pulse position within the stimulation pattern and the neuron (shown for Neuron#5 and Neuron#11). Five superimposed raw data traces are shown for each neuron for the responses to stimulation of ch#3 at various time points (as indicated by pu# and arrows below traces) within stimulation pattern S20. **(C)** Average EPSP amplitude, indicated as multiples of the average EPSP amplitude to the isolated single-pulse stimulation of the corresponding channel in the same neuron (Figure 6), shown for each neuron and each analysed within-pattern stimulation pulse (N=52), corresponding to a total of 56762 responses. For each analysed stimulation pulse, the stimulus pattern and the sequential position of the pulse within that pattern (pu#) for each respective stimulation channel (ch#) are indicated at the bottom (organised according to their order of occurrence within the stimulation patterns). White entries indicate that the response fraction did not surpass the spontaneous level of EPSP events by more than two standard deviations according to the automatic EPSP detection method and the response was hence discarded from further analysis. (**D**) Similar display as in A, but for the relative response latency time, i.e. the latency expressed as multiples of the average response latency time to isolated single-pulse stimulation of the corresponding channel. (**E, F**) Pairwise comparisons of differences in EPSP amplitudes between all neurons. For two sample stimulation pulse positions (S10, ch#3, pu#6; and F5, ch#3, pu#4, respectively) we analysed whether the changes in relative EPSP amplitude (compared to the neuron’s isolated single-pulse response) differed between neurons. The p-values of the Wilcoxon rank sum test for each pairwise comparison of the raw EPSP-like responses are reported as a colour code in the matrix (N=56 comparisons).

Responses to the individual stimulation pulses were analysed both manually and automatically. The manual part of the analysis consisted in a visually guided definition of the onset latency, amplitude height and latency-to-peak using the tailor-made point-and-click user interface (as in Figure 6A-C). The automatic part consisted of a detection of EPSP-like events using tailored template matching routines – its sole purpose was to identify if EPSPs evoked by a particular stimulation pulse were so infrequent that they were at risk of not surpassing the spontaneous occurrence of similar EPSP events (in the recording times in between the presentation of the stimulation patterns), in which case they were to be excluded from further analysis. EPSP templates consisted of a series of 5-20 time-voltage thresholds with individually variable voltage variance and were defined manually for each neuron based on a large sample of EPSP-like events (≫100) occurring after in-pattern stimulation pulses. They were visually confirmed to not omit mid to large EPSP-like events (> 3 mV) that occurred spontaneously at randomly sampled time points throughout the duration of the recordings. The response fraction of a neuron to each repetition of a stimulation pulse was defined as the number of repetitions evoking an EPSP-like event, as judged by the automated EPSP identification in the time range 4 - 18 ms after the pulse onset, divided by the total number of repetitions of that pulse. The baseline activity of that same EPSP-like event was determined by counting its spontaneous activity in time bins of 14 ms width (i.e. the same width as the response time window) for 12 consecutive bins preceding the onset of the stimulation pattern. As each stimulation pattern as a rule was repeated 100 times, we typically obtained a total of 1200 such bins. The response fraction of the spontaneous activity was obtained by taking the average activity across these 1200 bins. The threshold activity for the EPSP template, i.e. the response fraction that an in-pattern stimulation pulse needed to exceed in order to qualify as an evoked rather than spontaneous response, was defined as the mean plus two standard deviations of the response fraction of the spontaneous activity. If the response fraction was below the threshold activity, or if there was less than five manually detected EPSPs, the response of that in-pattern stimulation pulse was considered not significant and was discarded from further analysis.

#### Statistical analysis summary

For statistical evaluation of the identified response types, we used the Kruskal-Wallis test (Figure 2D-E). For pairwise tests of response fractions occurring under desynchronised brain states (Figure 5), we used paired t-test. For pairwise comparisons of EPSP-like responses (Figure 6–7), we used the Wilcoxon rank sum test for pairwise comparisons as the data was not obviously following a normal distribution.

### Data and software availability

Raw intracellular recording data can be accessed at FigShare (link on acceptance).

## Results

### Large variations in neocortical internal states and responses

While providing specific spatiotemporal tactile afferent activation patterns to the skin (Figure 1A), we made intracellular, whole-cell patch clamp *in vivo* recordings from single neocortical pyramidal cells (putative layer III-V pyramidal neurons in the somatosensory cortex (S1), Figure 1B), three of which were morphologically verified to be layer III pyramidal cells (Figure 1C). The time-varying states of the subnetworks connected to the recorded neuron generated a rich background of spontaneous Up and Down states, mixed with episodes of intermediate states, against which the responses evoked by the sensor input patterns could often stand out as distinctly different (Figure 1D). Responses evoked by the same stimulation pattern appeared to be impacted by the preceding state, as reflected in the spontaneous activity (Figure 1D, top two traces). In addition, in some cases, the spontaneous activity could even resemble the responses evoked by the stimulation (Figure 1D, bottom trace), in agreement with previous observations (Berkes *et al.*, 2011).

### Distinct types of responses on repeated application of identical tactile input patterns

Given a constant input pattern (Figure 1A), the response to any element in the stimulation pattern might still be modulated by cortical state changes, internally generated and/or impacted by prior elements of the pattern. Indeed, repeated delivery of one specific stimulation pattern generated multiple different responses that on visual inspection seemed to sort into a few different categories, or types (Figure 2A). In order to explore this separability, we applied a clustering method that sorted responses on basis of their membrane potential at each sample time point. In the illustrated case, this method indicated that the majority of the responses evoked by this specific stimulation pattern were separable into six recurring response types (Figure 2B). For the same neuron, the other seven stimulation patterns had 3-6 response types each. Across all neurons recorded, the responses of the intracellular membrane potential evoked by each stimulation pattern were on average divisible into 3.8+/−1.3 response types (Table 1).

**Table 1.** Number of response types and the performance measures for the response type separation for each stimulation pattern separately, based on PCA and kNN analysis. Summarised data for all neurons displayed for each stimulation pattern (based on matrices of the type shown in Figure 3A), and as a Grand Average across all stimulation patterns.

| MEAN (STD) | Grand<br>Average | F5 | S5 | F10 | S10 | F20 | S20 | F<sub>INF</sub> | S<sub>inf</sub> |
| --- | --- | --- | --- | --- | --- | --- | --- | --- | --- |
| Identified response<br>types (N) | 3.8 (1.3) | 3.5<br>(1.0) | 3.9<br>(1.4) | 4.0<br>(1.4) | 4.0<br>(1.6) | 4.1<br>(1.5) | 3.5<br>(1.8) | 3.7<br>(2.2) | 3.6<br>(1.8) |
| Type separation<br>accuracy (%) | 60.9 (8.1) | 59.5<br>(7.5) | 62.6<br>(14.2) | 61.8<br>(9.0) | 61.5<br>(13.2) | 56.7<br>(14.2) | 61.9<br>(13.0) | 60.7<br>(10.3) | 62.7<br>(12.2) |
| Fraction<br>'Ungrouped' (%) | 37.5 (14.6) | 39.5<br>(12.9) | 34.8<br>(12.8) | 33.1<br>(18.2) | 34.1<br>(17.6) | 43.4<br>(15.9) | 38.2<br>(18.0) | 38.6<br>(15.3) | 37.9<br>(20.4) |
| F1-score,<br>'Ungrouped' (0-1) | .48 (.18) | .51<br>(.21) | .48<br>(.19) | .45<br>(.21) | .45<br>(.21) | .54<br>(.18) | .50<br>(.21) | .48<br>(.22) | .47<br>(.26) |

**Table 2.** Quantification of single-pulse responses in the population of neurons. Quantified response variability across the entire population of neurons using the coefficient of variation (CV) measure. We investigated responses to inputs from 4 channels for 12 neurons, and hence calculated the CV for N=48 stimulus presentations, each repeated 100 times (4 exceptions, see Methods). For the CV of the response onset latency time we subtracted a fixed 4 ms delay for the cuneo-thalamo-cortical pathway. The corresponding values are displayed on the second row. Altogether, the data indicated that the three measured response parameters had a large internal variability.

| MEAN +/- STD | Peak amplitude | Response onset latency time | Latency to peak |
| --- | --- | --- | --- |
| CV | 0.43+/-0.10 | 0.28+/-0.10 | 0.29+/-0.08 |
| Values | 7.7+/-4.8 mV | 11.1+/-3.1 ms | 9.8+/-5.6 ms |

For the clustering method we used, the set thresholds of variability of the membrane potential were naturally critical factors for the outcome. A parameter sensitivity analysis showed how different parameter settings affected the outcome in two different sample neurons (Figure 2C). The aim for the parameter settings was to have as few ungrouped responses and as many groups as possible. However, across the population of neurons there was no single setting that would maximise these two factors across the whole population of neurons (Figure 2C). As shown in Figure 2C, the parameter settings that were chosen were located in the center region of a relatively smooth landscape of possible outcomes.

We next investigated if the clustered response types were different from each other using the Kruskal-Wallis test. We compared the responses that belonged to each response type to the responses evoked by the same stimulation pattern in the same neuron but belonging to the other response types. Whereas the Kruskal-Wallis test results for individual response types could fluctuate across different sample time points (Figure 2D), it was clear that the probability of H0 was substantially lowered during the time window that the stimulation pattern was on (0-350 ms, approximately). When we instead plotted the fraction of the response types that were statistically different (at p<0.05) from the other responses evoked by the same respective stimulation pattern, this separability was true for more than 60% of the response types across every time point for the duration of the stimulation patterns (Figure 2E). When the stimulation patterns had stopped, however, the separability dropped relatively rapidly (within 100-150 ms) to chance levels (Figure 2E). Furthermore, when the response type labels were shuffled among the responses evoked by the same stimulation pattern in the same neuron, the separability collapsed to chance level (0.05; red trace in Figure 2E) for all time points.

As another independent verification method, we next used Principal Component Analysis (PCA) to find out to what extent the full duration of the response types were distinctly different from each other. This method provided a measure of the accuracy, or the distinctness of separation of the individual responses, as summarised in the confusion matrix for the sample stimulation pattern in this neuron (Figure 3A). The mean accuracy across the different response types is indicated in the diagonal, and was on average 56.7% for this stimulation pattern (including ungrouped responses; the chance level was = 1/7 = 14.3%). Across the population of recorded cells, the accuracy of the separation of the responses into the identified response types was generally above 60% for each of the eight stimulation patterns (Table 1), verifying that the identified response types were composed of a set of responses that to a relatively high degree were orthogonal to each other. A sensitivity analysis of the parameter choice for the clustering method showed that the chosen parameter values did not provide the highest possible decoding accuracy (Figure 3B), but was instead a balance between this metric, the number of ungrouped responses and the number of response types identified (Figure 2C).

Whereas the majority of the evoked responses could be classified as belonging to one of the response types (Table 1), ‘ungrouped’ responses (Figure 3A) were by definition a much broader and heterogeneous class and consequently had a much greater risk of confusion (i.e. a low value in the diagonal and a higher prevalence of well-above-zero values outside the diagonal, Figure 3A). To evaluate the ‘ungrouped’ responses we used the F1 score, where a high value indicates a small risk of confusion with any of the specific classes of responses. In Figure 3A, the F1 score for ungrouped responses was 0.32, which does suggest some degree of confusion, even though the confusion matrix indicates that there wasn’t any specific response type that the ungrouped responses were confused with. Across the dataset, the F1 score was 0.48 (Table 1), indicating that across the data set, the ungrouped responses were overall well separated from the responses classified as belonging to the defined response types, and therefore potentially represented additional response types that our method could not reliably identify due to the limited number of repetitions (100 or less) available for each input pattern.

We also used PCA to compare the response types evoked by all eight stimulation patterns (Figure 3C). As each of the eight stimulation patterns on average evoked about four responses types, a large number of comparisons were made in this investigation. In this case, the accuracy of the response types (ungrouped responses were excluded here) was on average 57.6%. This is a remarkably high specificity, considering that the chance level for this comparison, for the number of response types being compared, was only 2.9% (for 34 response types compared the chance level is 1/34). Across all cells recorded, this cross-stimulation pattern response type accuracy was 38.2+/−21.0%. Hence, in each individual neuron, a majority of the evoked responses for a single stimulation pattern could be divided into distinct types that were separable with high accuracy from responses not belonging to that type, whether they were evoked by the same or by other stimulation patterns. In order to confirm that the reported decoding accuracy was not due to chance, we shuffled the stimulation pattern labels between the responses recorded in the same neurons. In this case, the decoding accuracy dropped substantially (Figure 3D), where the fact that it did not drop all the way to chance level was likely due to the circumstance that many of the evoked responses, just like the spontaneous activity (Figure 1D), did share some common features, owing to the physiological structure of the specific subnetworks that provided the input to each recorded neuron.

### Uniqueness of the response types between neurons

We previously found that the responses evoked by a given stimulation pattern were different between neurons (Oddo *et al.*, 2017), which on average was the case also in the present set of recordings (Figure 4A). Here, the issue was instead if the various response types evoked by the same stimulation pattern were distinct from each other across cells, which would indicate that the subnetworks connected to each neuron differed across the population of recorded neurons. Figure 4B illustrates a confusion matrix of a sample stimulation pattern, for which the accuracy was 42.7% (chance = 1/48 =2.1%) across the thirteen neurons (some neurons were recorded in the same experiment, as indicated). Across all eight stimulation patterns, this accuracy was 39.5+/−3.5%. Moreover, in the sample illustration (Figure 4B), we found that 45 out of 48, or 93.8%, of the response types were separable from the responses of all other types (i.e. decoding accuracy higher than chance). Across all stimulation patterns, 44.1+/−2.9 response types, or 89.6+/−3.0% of each response type identified, were similarly separable from all other response types evoked by the same stimulation pattern.

### Response types were not associated with specific ECoG states

Given that anaesthesia tends to increase the duration and depth of synchronised cortical activity states (Constantinople & Bruno, 2011), where responses could possibly become more stereotyped, one suspicion could be that some response types occurred only during synchronised cortical activity whereas other response types occurred only in the desynchronised state. However, comparing the prevailing ECoG state between individual responses classified to the same response type clearly demonstrated that this was not a general rule (Figure 5A). A response evoked in the middle of a desynchronised state could be classified to the same response type (Type 0) as another response evoked in the middle of a synchronised state, even as the latter response was even evoked in the middle of a starting Up state in the neuron (Figure 5A). To further show that a rising Up state in itself was not a determinant of the response type, a sample trace of another response type (Type 1) is illustrated in Figure 5B, alongside another trace also belonging to Type 1. We next systematically compared the distribution of the relative probability of the desynchronised ECoG state for each stimulation pattern (a total of 8 times 13 neurons = 104 data points) with the corresponding distribution for each of the 494 response types identified (Figure 5C, D). In Figure 5C, we only analysed the distribution for response types with more than nine individual responses/members. In Figure 5D, we also included response types with fewer than ten members, which resulted in a separate small peak at 0.0 and a few very high values for the type-separated responses. These outlier bars could be explained by chance omission or addition of one response in the desynchronised state, which could greatly impact the ratio for response types with fewer than ten members as the overall probability of being in the desynchronised ECoG state was around 20%. Nevertheless, under both conditions, the response types were not distributed significantly differently from the overall responses evoked by the same stimulation pattern according to the paired t-test (p=0.76 and p=0.68, respectively). Hence, the ECoG indicator of the global brain state was not predictive of the response type, suggesting that the response types were not an effect merely of this global brain state but rather the result of an interplay between the input and the dynamics of subnetwork-specific internal state evolution as the stimulus presentation unfolded.

### Neuron specific responses to the individual pulses that composed the stimulation patterns

We also performed a separate analysis of the responses evoked by the individual stimulation pulses, of which each stimulation pattern was composed. A striking feature of the responses to isolated single-pulse stimulations, delivered in between the stimulation patterns (analysed in Figures 1–5), was their highly variable nature (Figures 6A-C, Table 2), in contrast to the known consistency of the primary sensory afferent activation of this type of electrocutaneous stimulation (Bengtsson *et al.*, 2013). A statistical analysis moreover suggested that there was no order-dependence of the amplitudes for the responses evoked by the isolated single-pulse stimulations when delivered in chunks of five pulses separated by 0.3 ms (Figure 6D-E). In contrast, for the same single-pulse stimulations, when part of the stimulation patterns, the responses displayed a relative specificity that depended on the position of the pulse within the stimulation pattern (Figure 7), and which moreover differed between neurons (Figure 7A, B). Systematic analysis indicated that each neuron generated a unique pattern within the ‘mosaic’ of response parameter variations across the whole set of ‘within-pattern’ stimulation pulses (Figure 7C, D). This was quantified using a pairwise statistical comparison across all neurons, illustrated in Figure 7E, F for two sample within-pattern stimulation pulses. Across all 52 stimulation pulse positions tested (see Methods), statistically significant differences between the neurons (at p<0.05, Wilcoxon rank sum test for pairwise comparisons; the analysis only included responses that were non-white entries, i.e. statistically significant, in Figure 7C, D) were found in 65% of the comparisons of the amplitudes, 78% of the comparisons of the time-to-peaks and 86% of the comparisons of the response latency times. Thus, for the majority of the within-pattern stimulation pulses, the changes in response amplitude, time-to-peak and response latency relative to the isolated single-pulse stimulation were statistically different between the neurons, corroborating the results from Figure 4B that each neuron responded with partly unique time-voltage curves across the duration of the stimulation patterns. Altogether, the results in Figure 7 show that the outcome of the response type identification analysis cannot be explained by a generic sequence of paired-pulse depression or facilitation phenomena, since these would affect all neurons in the same way.

## Discussion

Our results show that the responses evoked by tactile sensory input patterns fall into a limited subset of preferred response states that are specific to each input pattern and each cortical neuron. The relative orthogonality of these recurring response types (Figure 3A, Table 1), the identification of which was made possible by the high resolution of the recording and precise repeatability of the stimulation method, indicates that they result from specific response states of the cortical networks, rather than being noise or specific patterns of spontaneous activity unrelated to the stimuli. The finding that the response types were relatively unique even when all the response types evoked across all the eight stimulation patterns were compared (Figure 3C) is another strong piece of evidence against that possibility. Moreover, the observation that the response types are relatively unique for each neuron (Figure 4B) indicates the existence of neuron-specific subnetworks, which could reflect specific aspects of the current global internal brain state (Figure 8). This is conceivable since the global internal states subsampled by each neuron, in turn, are likely to encompass large parts of the neocortex (Spanne & Jorntell, 2015; Enander *et al.*, 2019; Stringer *et al.*, 2019a; Stringer *et al.*, 2019b; Wahlbom *et al.*, 2019). The existence and regulation of such internal brain states can be expected to be essential for forming the percepts (or illusions), generated by given sensor activation patterns (Geldard & Sherrick, 1972; Robles-De-La-Torre & Hayward, 2001). Our results indicate that the neocortical network physiology has a non-uniform, non-continuous landscape of solutions that is adaptable to the time-evolving match between the subnetworks’ internal states and the actual sensory activation pattern received.

**Figure 8.**
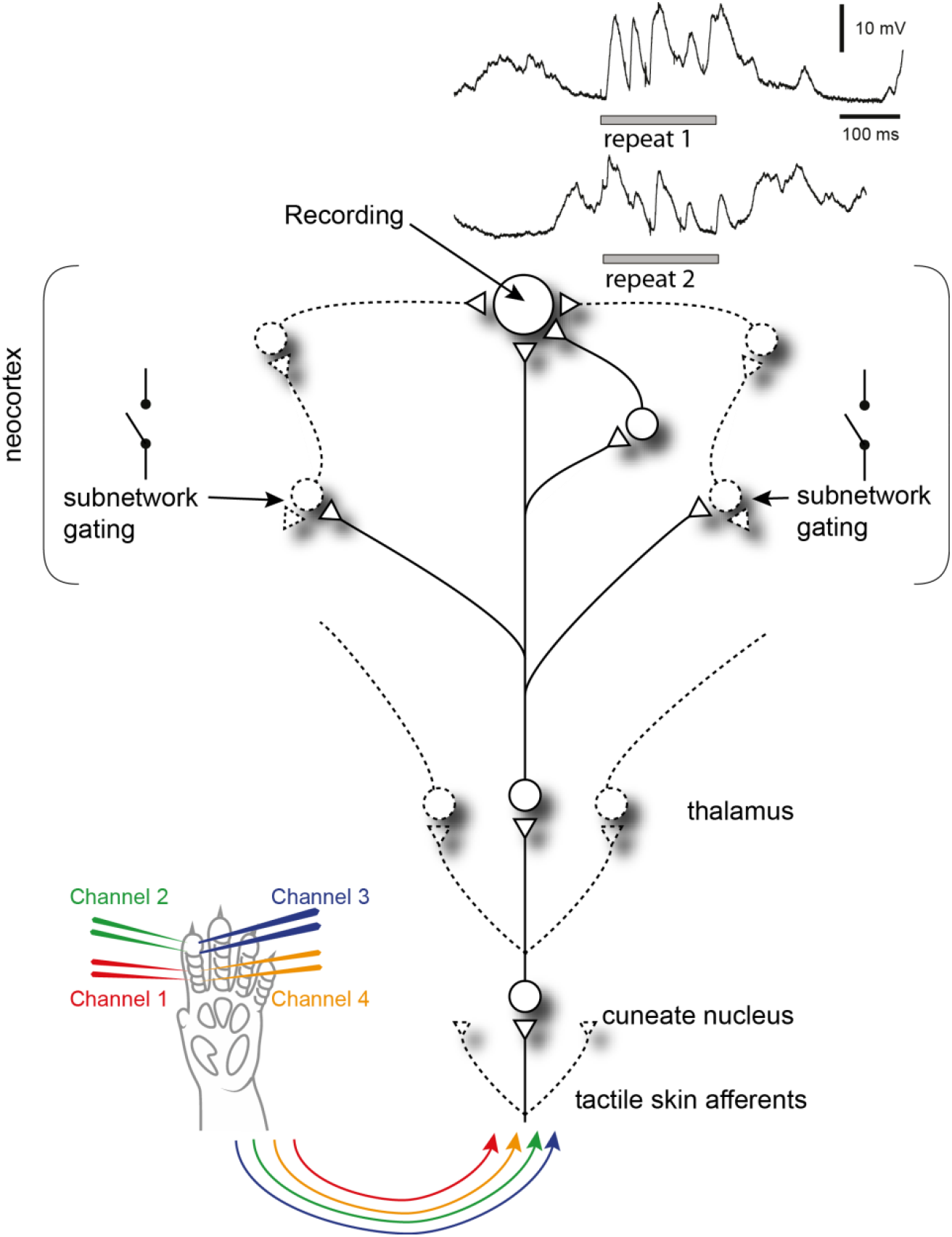
Summary of findings. Recorded neocortical neurons are provided by a high number of synaptic inputs, each of which could potentially be controlled by a different subnetwork. Each subnetwork may be gated by other, relatively independent, events in the global neocortical circuitry. When tactile afferent stimulation is provided at different points in time, these subnetworks may have different initial states, which will impact which subnetworks are gated in or out, which in turn will affect the appearance of the intracellular response (two traces shown in the top right corner). Circles are neurons, triangles are synapses, lines are axons and variable connections are drawn by dashed lines.

### Methodological considerations

Our response types should not be regarded as fixed entities, as for example the specific types detected could grow or shrink with altered parameter settings of the clustering method (see Figure 2C) and there was a degree of response variability within each identified response type (Figure 2B). However, as each response type could subsequently be demonstrated to have a degree of orthogonality to the responses belonging to other types (Figure 3A, Table 1), our data show that the dynamics of the neocortical network created a multitude of alternative, non-continuous response states to any given sensory input pattern. It is highly unlikely that our identified average number of four response types per stimulation pattern represents anything near an actual upper limit – if we had had the possibility to repeat each stimulation pattern millions of times, rather than one hundred, a higher number of response types would most likely have been possible to identify. Nevertheless, as shown in our sensitivity analysis, where we tested hundreds of different settings of our clustering algorithm, almost every clustering setting with more than one identified response type indicated that the identified clusters were objectively separable using PCA+kNN (Figure 3B). Hence, our analysis indicates the existence of discontinuity, or local minima, in the response states of the cortical network.

Alternative approaches to clustering, like machine learning networks or k-means, were not explored here, as these approaches commonly require a target aim of how many clusters should be sought for. Our clustering method was instead designed to objectively identify how many clusters could be observed, based on clearly defined metrics of the neurophysiological recording data.

How many possible response types exist and what specific membrane voltage curves they would create is more than likely to depend on the condition, and it would not be surprising if they are instantiated differently in the awake behaving animal, where they may be varying even further with the ‘state of mind’, and differently still when there is an accurate prediction of an expected sensory input being made. However, apart from being methodologically nearly impossible to explore in the awake animal (due to the requirement of achieving identical stimuli over a long time), a correct analysis of such data in this respect would have required a precise estimate of what the experimental subject is thinking at the time of the stimulus delivery, which is today not theoretically possible.

To what extent could our results be a ‘product’ of the anaesthetics? Anaesthesia increases the probability of the thalamocortical system to enter episodic, coordinated oscillatory modes (Amzica & Steriade, 1998; Constantinople & Bruno, 2011), or synchronised states, even though they are certainly not uncommon in the awake state (Petersen *et al.*, 2003; Poulet & Petersen, 2008; Bennett *et al.*, 2013) (synchronised states are widely observed also in awake humans, (Sachdev *et al.*, 2015)). However, in between such episodes, there are periods of desynchronised activity, or states, that do not obviously differ between the awake and the anaesthetised animal, though anaesthesia could be expected to lead to an overall activity reduction within desynchronised states (Constantinople & Bruno, 2011). Importantly, the response types were not specifically associated with any of these two opposite global states (Figure 5), which is a strong argument against the possibility that the response types are a peculiarity created by the increased time spent in synchronised state that is a consequence of the anaesthesia. Hence, whereas our recordings do not reflect natural integrated thought processes, they can still provide important information about underlying governing principles of information processing, principles which would be much harder to disentangle in awake recordings. However, data from awake behaving animals are compatible with the presence of multi-structure cortical states as indicated by the fact that the activity of a population of neocortical neurons can follow different trajectories of interdependencies depending on the motor task performed (Churchland *et al.*, 2012; Gallego *et al.*, 2017; Russo *et al.*, 2020).

### Relation to previous literature

Previous analyses of state-conditioned intracellular signal variations evoked by somatosensory inputs have largely focused on the binary issue of Up and Down states using single-shot or brief inputs. The seemingly paradoxical finding that inputs provided in an excited, Up, state result in much lower responses than the same input in a Down state (Petersen *et al.*, 2003) was in a detailed conductance analysis indicated to be partly due the shunting effect caused by the Up state being associated with a high level of excitatory synaptic input but also in parallel a high level of inhibitory synaptic input (Hasenstaub *et al.*, 2007). Moreover, it was observed that stimuli during Down state could often evoke disproportionately large responses, by inducing transitions to Up state. More recent studies suggest such effects to be inducible due to that the neocortex is a system with criticality phenomena, where small changes at critical states can cause avalanche effects of increased excitability that spread widely through the cortical network (Wright & Wessel, 2017; Johnson *et al.*, 2019). Although our data indicate a much more fine-grained subdivision of cortical states than the binary dichotomy between Up and Down states, such avalanche effects could be the underlying reason for the response types we observed here. If each subnetwork for example exhibits partly independent criticality, where each individual synaptic input to a neuron could in theory represent the activity state of partly independent subnetworks, this would be compatible with the fact that different neurons exhibited different response types to the same inputs. Moreover, the data suggest that different individual neurons are connected to unique combinations of such subnetworks (Figure 8).

### Implications for our view on the neocortical mode of operation

Although our data naturally do not allow an identification of the full perceptual processes, they explore the physiological brain mechanisms that support such processes, which likely corresponds to mechanistic underpinnings of expectations (Loeb & Fishel, 2014). The findings suggest that the multidimensional latent state defined by the large populations of neurons across the neocortex (Stringer *et al.*, 2019b) works according to attractor-like dynamics (Ringach, 2009) with multiple attractive states for each given input. The interaction between the input pattern and the cortical state at the moment of stimulus onset would thus cause the cortical network to fall into one specific out of many possible attractive states. Since different neurons would be coupled to different subnetworks of this global network, the same attractive state would potentially impact different neurons in different ways, as our data indicate (Figure 4B). The specificity of the response types observed in individual neurons would thus be local subnetwork-instantiations of the time-evolving input-updated brain-wide state estimations of world and body, hence fundamental mechanisms for forming rich perception, and illusions. We expect these principles to reflect a general computational strategy used by the neocortex across all sensory systems.

## Acknowledgments

The authors thank Jerry Loeb (USC Los Angeles) and Matthias Kohler (TU Munich) for pre-reviewing our manuscript. Funding: This work was supported by the EU Grant FET 829186 ph‐coding (Predictive Haptic COding Devices In Next Generation interfaces), the Swedish Research Council (project grant no. K2014-63X-14780-12-3).

## Author Contributions

J.N. and H.J. planned the study and conducted the experiments. J.N., J.M.D.E. and H.J. designed the analysis. J.N., J.M.D.E. and H.M. conducted the analysis. J.N. and H.J. wrote the paper.

The authors declare no competing interests.

## Notes

### Competing Interest Statement

The authors have declared no competing interest.

